# Tumor susceptibility gene-101 regulates glucocorticoid receptor through disorder-mediated allostery

**DOI:** 10.1101/2020.02.02.931485

**Authors:** Jordan T. White, James Rives, Marla E. Tharp, James O. Wrabl, E. Brad Thompson, Vincent J. Hilser

**Affiliations:** Department of Biology at Johns Hopkins University, Baltimore, MD 21218; Department of Chemistry at Johns Hopkins University; Sealy Center for Structural Biology and Molecular Biophysics in the Department of Biochemistry and Molecular Biology at Univ. of Texas Medical Branch, Galveston, TX; T. C. Jenkins Department of Biophysics at Johns Hopkins University

**Keywords:** glucocorticoid receptor, tumor susceptibility gene-101, allostery, allosteric regulation, transcription, transcription coregulator, transcription regulation, thermodynamics, biophysics

## Abstract

Tumor Susceptibility Gene-101 (TSG101) is involved in endosomal maturation and has been implicated in the transcriptional regulation of several steroid hormone receptors (SHRs), although a detailed characterization of such regulation has yet to be conducted. Here we directly measure binding of TSG101 to one SHR, glucocorticoid receptor (GR). Using biophysical and cellular assays, we show that the coiled-coil domain of TSG101; 1) binds and folds the disordered N-terminal domain (NTD) of GR, 2) upon binding, improves DNA-binding of GR *in vitro*, and 3) enhances the transcriptional activity of GR *in vivo*. Our findings suggest that TSG101 is a *bona fide* transcriptional co-regulator of GR.

## Introduction

Transcription factors (TFs) are multi-domain, allosteric proteins, in which the binding of one effector ligand to its specific domain causes a conformational change in one or more other domains, so as to alter transcription. Most TFs contain intrinsically disordered regions (IDRs), which though critical for TF function, are still not well understood (1). Generally, TF IDRs consist of large ensembles of rapidly interconverting conformers, including in many cases, conformers that contain some degree of structure. By definition then, allostery arises due to an effector-driven shift in the distribution of states such that states that are capable of initiating transcription are either promoted or repressed, thus affecting transcriptional activity accordingly (2,3). We have adopted the glucocorticoid receptor (GR) as a model system for our studies because the NTD of GR is known to be intrinsically disordered (ID) and also to be critical for the TF function of the receptor.

The GR is a 3-domain protein comprised of (from N to C terminus): N-terminal (NTD), DNA-binding (DBD), and ligand-binding domains (LBD), where L in this case refers to steroid or steroid substitute. However, if the LBD is deleted, the 2-domain NTD-DBD is constitutively active due to a potent activation function-1 (AF1) region located in its NTD, a fact that makes the GR NTD-DBD amenable to exploring the allosteric impact of DNA binding on transcriptional activation, as well as the effects of changes to the NTD on DNA binding (Fig. 1A). We and others have shown that the ID NTD can be made to cooperatively fold into conformers containing both 2° and 3° structure, by use of; 1) protective organic osmolytes (4–7), 2) known binding partner proteins (8–10), 3) phosphorylation of key serines (11,12), and 4) addition of the DBD (13,14).

**Figure 1.**
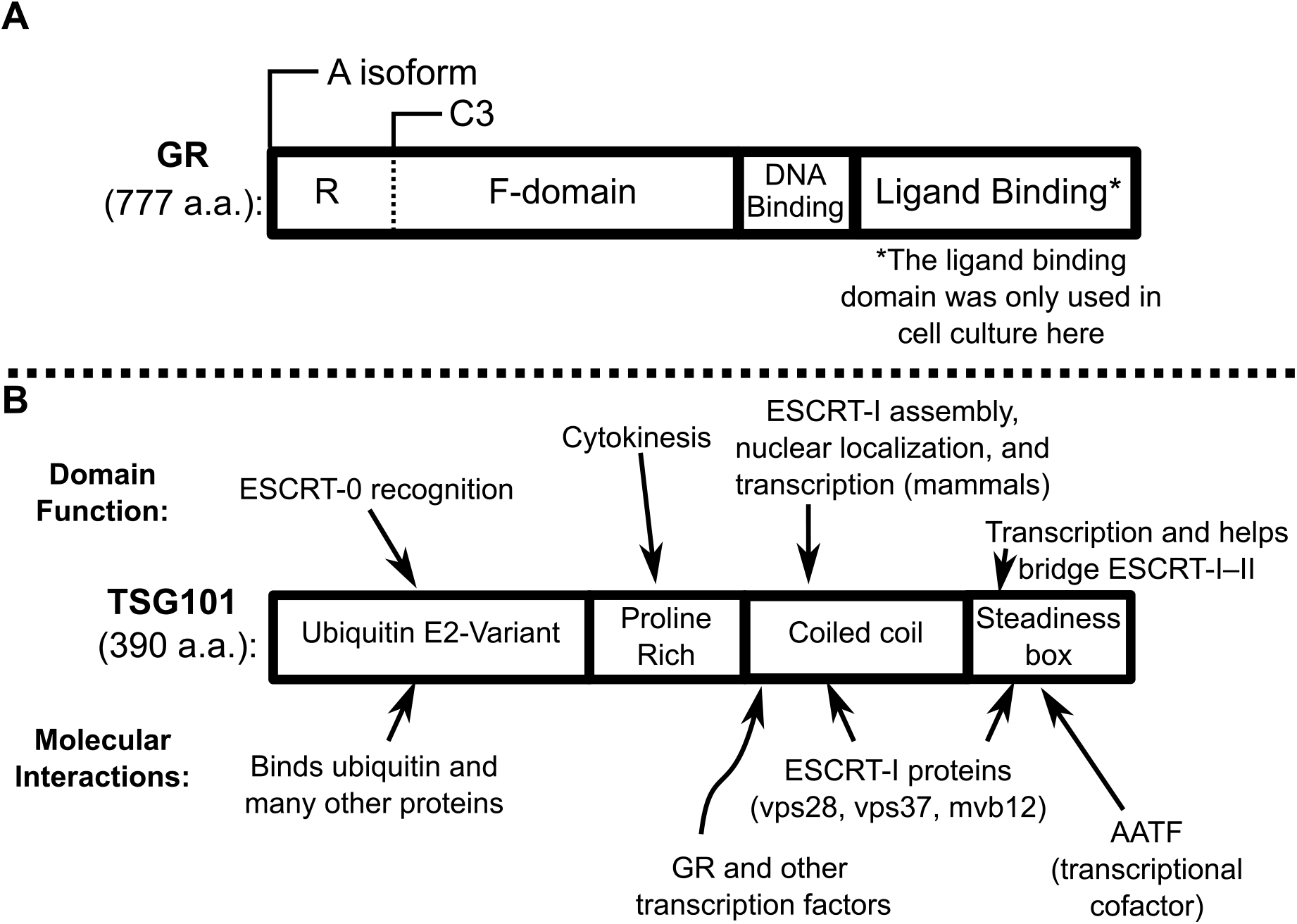
The domains and functions of GR and TSG101. Domains are rectangles oriented from N- to C-terminus, scaled with respect to size. **A)** Two of GR’s translational isoforms are indicated above the GR diagram. The A NTD isoform contains a regulatory (R) domain that alters the stability and transcriptional activity of the GR (13). The functional domain (F) binds transcriptional cofactors and is also called activation function 1 (AF1) in the literature. AF1 has been shown to bind: TSG101, Med14, TBP, CBP, Ada2, and many other cofactors (7,23,46,47). **B)** TSG101 domain functions are listed at the top, and the coiled-coil’s nuclear involvement is highlighted. Some examples of protein binding partners are listed at the bottom, see references for a more complete listing (20–23,25,48–54).

Previously, we hypothesized, using a rigorous thermodynamic model, that under a wide range of initial energetic conditions, inclusion of an IDR in a protein could enhance its allosteric behavior (15). Specifically, in a 2-domain model in which each domain is capable of binding its own specific ligand, when one or both domains is ID, binding of ligand to one domain can greatly influence the probability of the second domain being in a favorable folded state for binding its ligand. Further, we showed this ensemble allosteric model (EAM) could be applied to proteins with multiple domains, allowing us to probe the role of ID in mediating allosteric control in numerous domain topologies (16). Indeed, in validating this model, we demonstrated that DNA binding affinity and transcriptional activity for a series of isoforms of GR could be described in the context of inter-domain coupling in the GR (13), thus, establishing one tenet of the EAM, that stabilization of an input domain (i.e. the DBD) could affect the function of the activation domain.

Here, we set out to test the reciprocal prediction of the model; that binding a transcriptional cofactor ligand to the NTD could either positively or negatively (depending on the sign of the coupling) tune the affinity of the DBD. To address this question, we utilized the 2-domain GR system described above (13). Importantly, within this construct, the DBD is known to be a structured, albeit highly dynamic, globular protein, which adopts the canonical zinc-finger binding motif (17,18). Conversely, we showed that although the NTD of GR appears in its entirety to be an IDR, it actually consists of two adjacent, but functionally distinct IDRs. The first IDR, which we call the functional (or F-domain) includes residues 99-420. We showed that the F-domain exists as an equilibrium between a disordered (or unfolded) state and a folded ordered state, and that the folded state of the F-domain is responsible for transcriptional activation.

The second functionally distinct IDR of the NTD consists of residues 1-98 (Fig. 1A), and this region serves as a variable-length regulator (or R-domain). The R-domain influences both the F-domain of the NTD as well as the DBD, providing GR with genetically “tunable” energetic frustration (13), whereby DNA binding to the DBD both directly stabilizes (and thus activates) the F-domain, but also indirectly destabilizes (therefore, repressing) the F-domain through stabilization of the R-domain. Thus, our results revealed that all the GR NTD-DBD isoforms that contain the AF1 region (e.g., A, B1, B3 C1, and C2), with the exception of the C3 isoform, actually consist of three domains; the DBD, the F-domain, and variable lengths of the R-domain. Because the GR C3 is the only isoform lacking the R-domain (19), it represents an ideal system to interrogate the direct impact of changes to the NTD on the DBD, as the absence of any R-domain should greatly simplify interpretation.

As a transcriptional cofactor for these studies we selected the Tumor Susceptibility Gene 101 (TSG101), a protein that is primarily known for its role in the ESCRT-I complex (i.e., Endosomal Sorting Complex Required for Transport-I), promoting endosomal maturation (19,20). However, it has also been reported to bind to the GR, and to possibly act as a cofactor for GR and other TF’s (Table S1 and (21–25)). The domains of TSG101 that are critical to these interactions are shown in Figure 1B. Herein, we investigate the binding of TSG101 to GR and the ability of this binding to allosterically affect GR DBD binding to its canonical palindromic DNA binding site (i.e., the GR response element (GRE)), as well as influence subsequent transcriptional activity.

## Results

### TSG101 binds within the GR NTD Activation Function-1 (AF1) region

It has been reported previously that the coiled-coil region of TSG101 (TSG101cc) binds to the AF1 region of the GR NTD (23). Here we set out to establish whether this binding is specific and whether it affects the functional coupling we previously revealed in GR (13). Importantly, we confirmed previous results and extended our understanding using quantitative binding studies and protease protection assays. First, we examined the binding of TSG101cc to the single-domain GR C3 NTD and to the two-domain GR C3 NTD-DBD (Fig. 2). Our results reveal that TSG101cc binds to the two constructs with 1:1 stoichiometry, with TSG101cc binding more tightly to the isolated C3 NTD (K_d_ = 19.6 ± 1.0 μM, see Table 1) than to the two-domain C3 NTD-DBD (K_d_ = 72.6 ± 14.6 μM). Similar results were obtained with the GR A isoform (Fig. S1), and importantly, TSG101cc does not bind the isolated DBD (data not shown).

**Figure 2.**
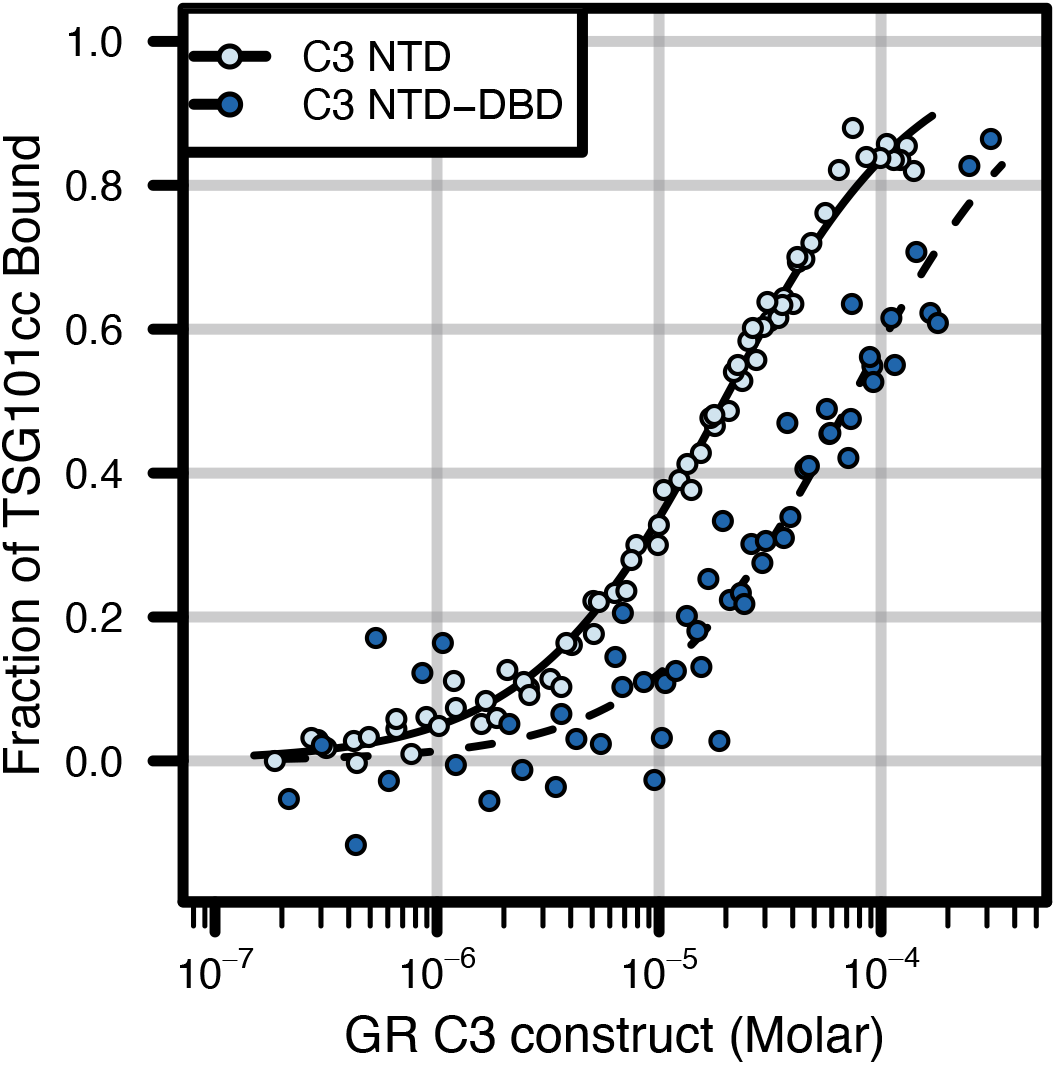
Quantitative binding of TSG101cc to C3 constructs of GR. Data from fluorescence anisotropy, globally fitted and converted to fraction of TSG101cc bound. Fluorescence anisotropy of pyrene labeled TSG101cc was followed while titrating either the GR C3 NTD (pale blue circles) or C3 NTD-DBD (dark blue circles). The data were fit to a single-site binding model (K_d_’s: C3 NTD = 19.6 ± 1.0 μM, C3 NTD-DBD = 72.6 ± 14.6 μM). See Supporting Information for details of fitting and Figure S1 for similar data on the GR A isoform.

**Table 1.** Globally fitted binding constants. Apparent binding constants are shown, all collected in the context of our fluorescence anisotropy buffer (methods). The first four rows refer to binding of GR constructs and TSG101cc in Fig. 2 and Fig. S1, and the rest of the table refers to binding of GR NTD-DBD and DNA in Figure 5 and S7. Parentheses next to the fitted K_d_’s indicate the number of replicate data sets that were globally fit. Errors are ± 95% confidence intervals of the non-linear least squares fits. 6-FAM construct acronyms: Pal = palindromic consensus, GilZ = GRE from glucocorticoid induced leucine zipper, Tat = tyrosine amino-transferase derived (45). For details of GREs used, see Supporting Information and Fig. S7.

| **GR** | **TSG101cc** | **6FAM-GRE** | **K<sub>d</sub>** |
| --- | --- | --- | --- |
| A NTD | + pyrene | N/A | 20.9 ± 9.4 μM (3) |
| C3 NTD | + pyrene | N/A | 19.6 ± 2.0 μM (3) |
| A NTD-DBD | + pyrene | N/A | 33.6 ± 15.2 μM (3) |
| C3 NTD-DBD | + pyrene | N/A | 72.6 ± 14.6 μM (3) |
| A NTD-DBD | – | Pal | 212 ± 12 nM (4) |
| A NTD-DBD | + | Pal | 100.3 ± 8.0 nM (5) |
| A NTD-DBD | – | GilZ | 241 ± 19 nM (2) |
| A NTD-DBD | + | GilZ | 183 ± 23 nM (2) |
| A NTD-DBD | – | Tat | 287 ± 37 nM (2) |
| A NTD-DBD | + | Tat | 128 ± 43 nM (2) |
| C3 NTD-DBD | – | Pal | 229 ± 24 nM (3) |
| C3 NTD-DBD | + | Pal | 94.5 ± 5.1 nM (3) |
| C3 NTD-DBD | – | GilZ | 238 ± 26 nM (2) |
| C3 NTD-DBD | + | GilZ | 183 ± 19 nM (2) |
| C3 NTD-DBD | – | Tat | 298 ± 43 nM (2) |
| C3 NTD-DBD | + | Tat | 192 ± 55 nM (2) |

To determine the location on the GR that interacts with TSG101cc, we conducted limited proteolysis of the bound complex. As seen in the electrophoretic gels (Fig. 3A), trypsin digestion of GR C3 NTD in the presence TSG101cc resulted in protection of a series of 14.8-20.9 kDa peptides from the GR AF1 region. Mass spectroscopic analysis of the protected bands identified a 9.1 kDa fragment of GR (Fig. 3B, also see Figs. S2-3 and Table S2), leaving about 10 kDa unaccounted for (although see below). These results agree with previous yeast-two hybrid analysis (23), and suggest specific binding to the NTD, consistent with the observed binding effects (Fig. 2)

**Figure 3.**
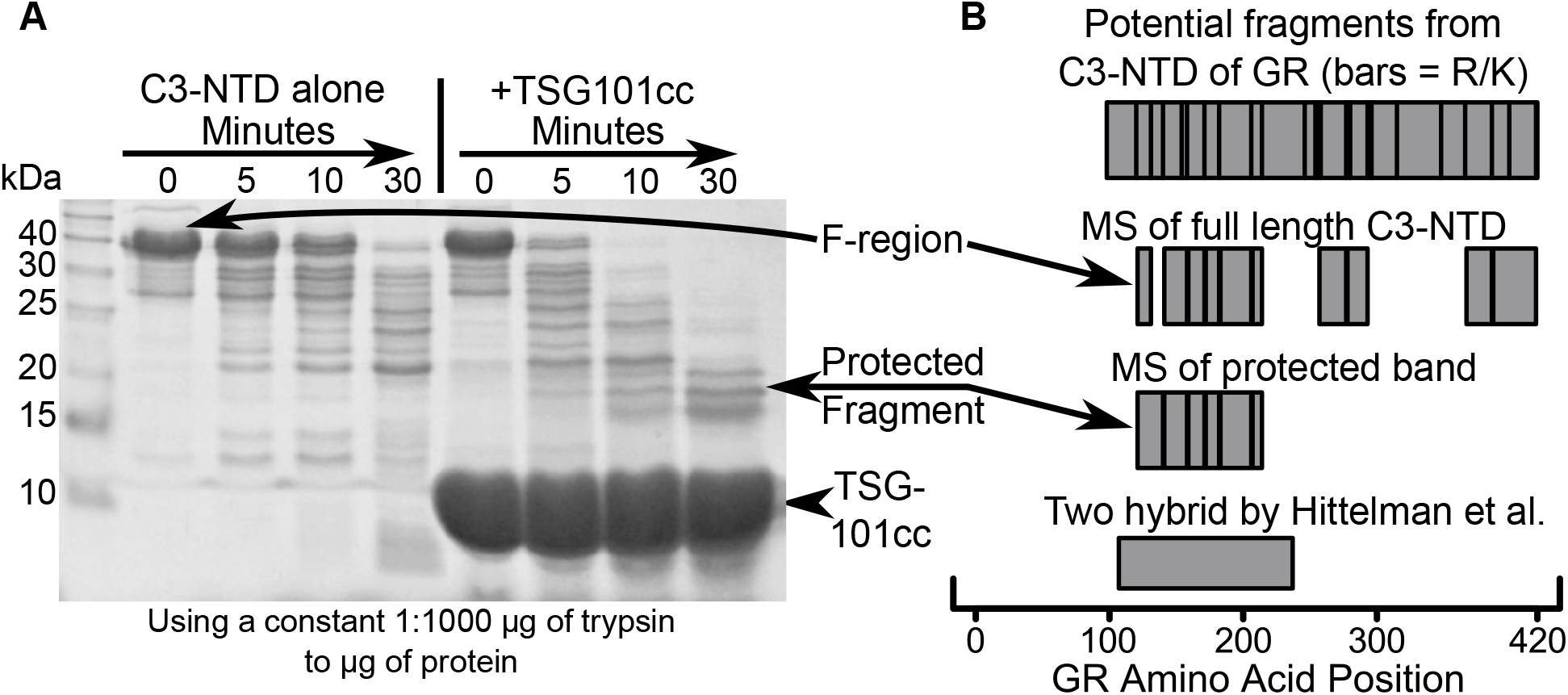
TSG101cc protects the N-terminus of GR from limited protease digestion. **A)** Polyacrylamide gel stained with Coomassie dye. Digestion of GR C3 NTD alone (left) produces numerous fragments. Addition of TSG101cc (right) causes formation of a protected fragment. Note that far more trypsin was added to the +TSG101cc reaction to maintain a constant ratio of trypsin:total protein. **B)** Illustrated results of MALDI mass spectrometry of bands cut from a gel, as shown in part A, and fully trypsinized overnight. Possible fragments are shown at the top, with trypsin cut sites (Arg or Lys) as black bars. In descending order, the next two sets of boxes represent the fragments actually observed for unprotected full-length GR C3 NTD and the protected band, respectively. The box at the bottom represents the sequence used as bait in the yeast two-hybrid experiment of Hittelman et al. (23). See Figures S2-3 and Table S2 for mass spectrometry, controls, and an example of GR A NTD digestion.

To gain insight into the structural organization of the bound TSG101cc:GR C3 NTD, the cross-linker disuccinimidyl tartrate (DST) was employed, resulting in cross-linking between TSG101cc and a ~19 kDa stretch of the GR C3 NTD (Fig. S4 and Tables S3-8). The results indicate that the GR binding site stretches from approximately residue 121 to 301. Of note, fluorescence anisotropy suggests that the AF1 core region (a.a. 187—244), which binds TATA-box Binding Protein (TBP (26)), does not appear to interact with TSG101cc (Fig. S5). This finding taken together with the protease protection and cross-linking results, indicates that the interactions with TSG101cc involve the amino acids prior to (i.e.~120-200) and following (i.e.~240-300) residues in the AF1 core. Interestingly, the orientation of the cross-links is suggestive of an inversion in the register of the interaction, whereby the GR NTD is aligned parallel with TSG101cc but then doubles back on TSG101cc, aligning in antiparallel (Fig. 4 inset). Importantly, these results are consistent with a model whereby the TSG101cc—which is known to form a heterotrimeric coiled-coil in its functional ESCRT complex (27)—interacts with the GR NTD in the two regions before and after the AF1 core residues (5), thus mimicking the trimeric coiled-coil seen in the ESCRT complex. We note that this proposed interaction model is also consistent with our structural thermodynamic analysis of GR using the COREX algorithm, whereby the two regions of the GR NTD that interact with TSG101cc are predicted to be highly helical (See Fig. 4, Supporting Information, Fig. S6, Supporting Excel Spreadsheet, and (28)).

**Figure 4.**
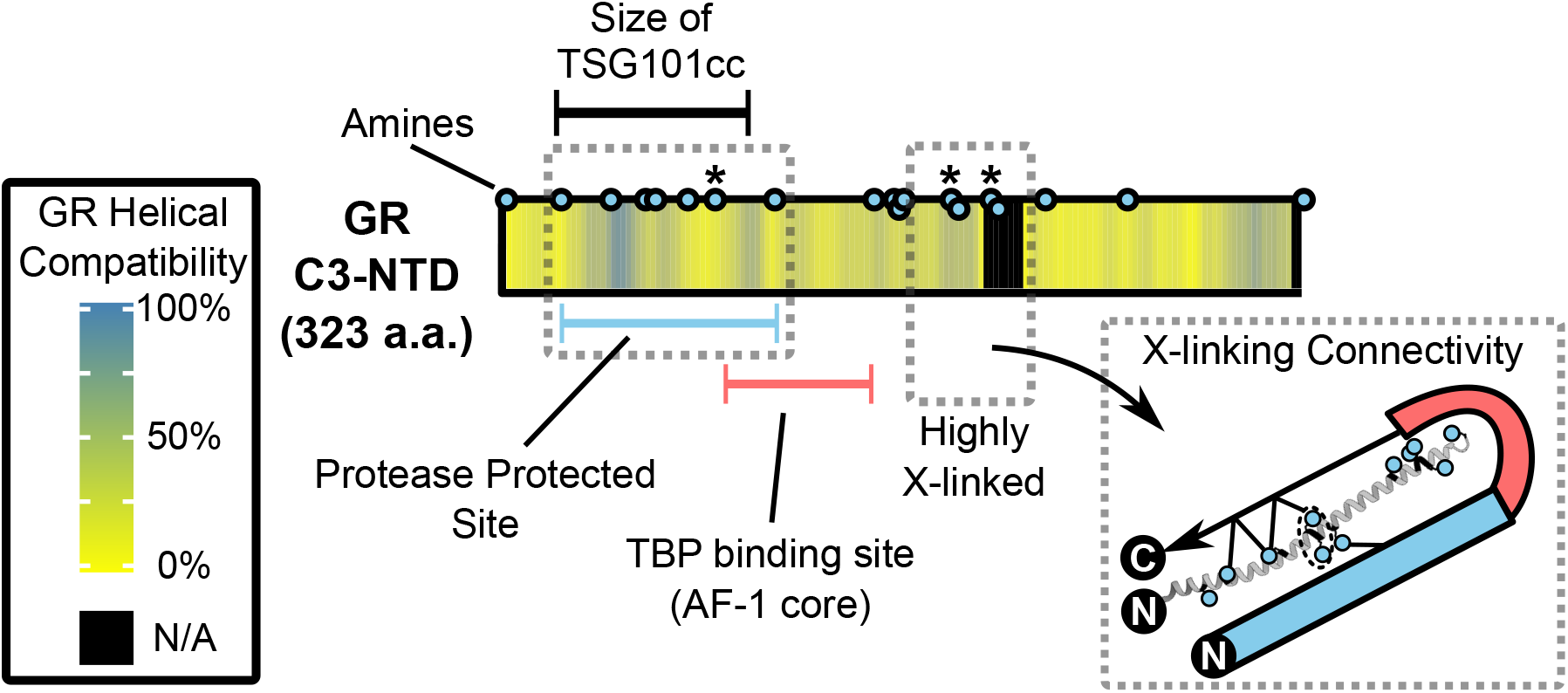
Cross-linking results and model for GR C3:TSG101cc binding. Representation of GR C3-NTD and TSG101cc cross-linking data. Amines within TSG101cc and GR C3 NTD are shown as blue dots (lysines or N-termini). An asterisk (*) indicates a cross-linked position on GR (raw data in Fig. S4 and Tables S3-8). The GR NTD sequence was also computationally analyzed for helical compatibility (28,35,55), represented by heat map coloring of the GR box in the center. “N/A” indicates a sequence around GR a.a. 300 that did not match any known structure and it is colored black. See Figure S6 and Supporting Excel file for the helical compatibility data. Around the GR box are three bars: the blue bar is in reference to Fig. 3, the red bar is the AF1 core of the literature (56), and the black bar represents the a.a. length of the TSG101cc. **Bottom Right Inset:** Model of the heterodimer, based on the following: 1) flipped cross-links, 2) helical compatibility of the GR sequence, 3) 1:1 binding of GR NTD and TSG101cc (Fig. 2 and Fig. S4), and 4) our finding that the GR AF1 core does not bind TSG101cc (Fig. S5). The TSG101 coil is in gray and the cross-linked lysines are shown as blue dots with lines going roughly to the appropriate site on GR. One cross-link was ambiguous because of adjacent lysines—indicated by a dashed circle. The GR protease protected site and TBP binding site are colored as in the main figure. The black spheres “N” and “C” indicate N and C-termini.

### TSG101cc alters GR NTD-DBD affinity for DNA sequences corresponding to the GRE

Previously, we presented an ensemble based thermodynamic model that allowed us to investigate the role of ID in mediating allostery (15,16). Using this model, we demonstrated the importance of ID in the allosteric behavior of 2 and 3-domain proteins (15,16). Applied to the GR, the model predicts that the DBD and the AF1-containing region of the NTD (termed the F-domain in (4)) are positively coupled, meaning stabilization of the high affinity state of the DBD increases the stability of the active conformation of the F-domain. If this model behaves reciprocally, binding of a ligand (e.g. TSG101cc) to the active state of the F-domain should also enhance the binding of the GR DBD to its ligand, the palindromic sequence of DNA that forms a GRE. Consistent with this prediction, the DNA binding affinity of the TSG101cc-bound GR C3 isoform is increased relative to the GR C3 alone, when tested on DNA sequences corresponding to several symmetric GREs, although the magnitude of the increase appears to be sequence dependent (Fig. 5, Table 1, and Fig. S7-8). Importantly, the relative increase in affinity of GR C3 NTD-DBD for DNA when bound to TSG101cc is also observed for the GR A NTD-DBD isoform (Fig. 5), suggesting that the presence of the repressor R-domain does not qualitatively affect the ability of TSG101cc to allosterically regulate the DBD through its interaction with the F-domain.

**Figure 5.**
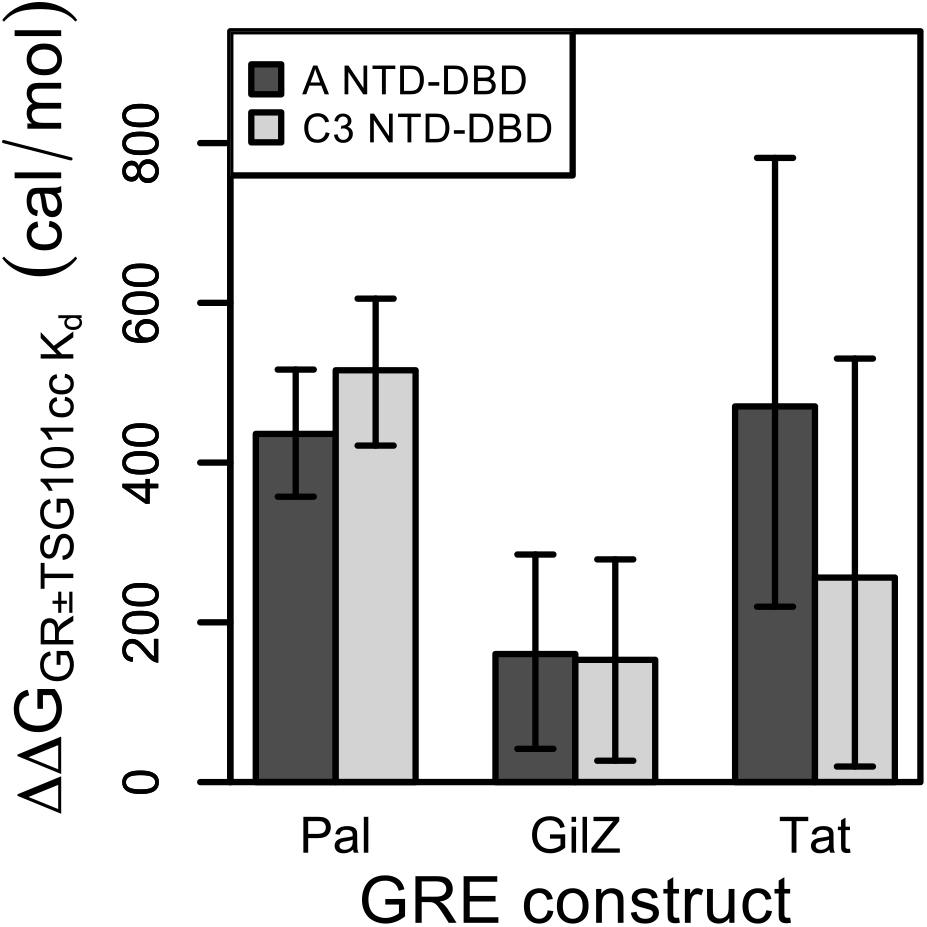
Binding of GR A/C3 NTD-DBD to GRE constructs with or without TSG101cc. The raw data were fit globally to extract K_d_’s, then converted to ΔΔG relative to the case without TSG101cc. Error bars are the additive 95% confidence intervals from two fits (K_d_ with versus without TSG101cc). See Table 1 for sample sizes and the fitted K_d_ values. See Fig. S7 for the fitted data. Pal = palindromic consensus, GilZ = GRE from glucocorticoid induced leucine zipper, Tat = tyrosine amino-transferase derived (45). See Supporting Information for the DNA sequences used.

### The effect of energetic frustration on TSG101:GR A binding

The above experiments reveal an overall positive allosteric coupling between TSG101cc binding to the GR NTD and DNA binding to the GR DBD. We also tested the reciprocal effect which would stipulate that DNA-bound GR should bind more tightly to TSG101cc. By titrating TSG101cc binding in a series of experiments with GR bound to palindromic GRE, we obtained an apparent binding affinity for TSG101cc and DNA-bound GR (Fig. 6A-B). The results confirmed the prediction: the apparent TSG101cc binding affinity is more than ten-fold tighter in this context.

**Figure 6.**
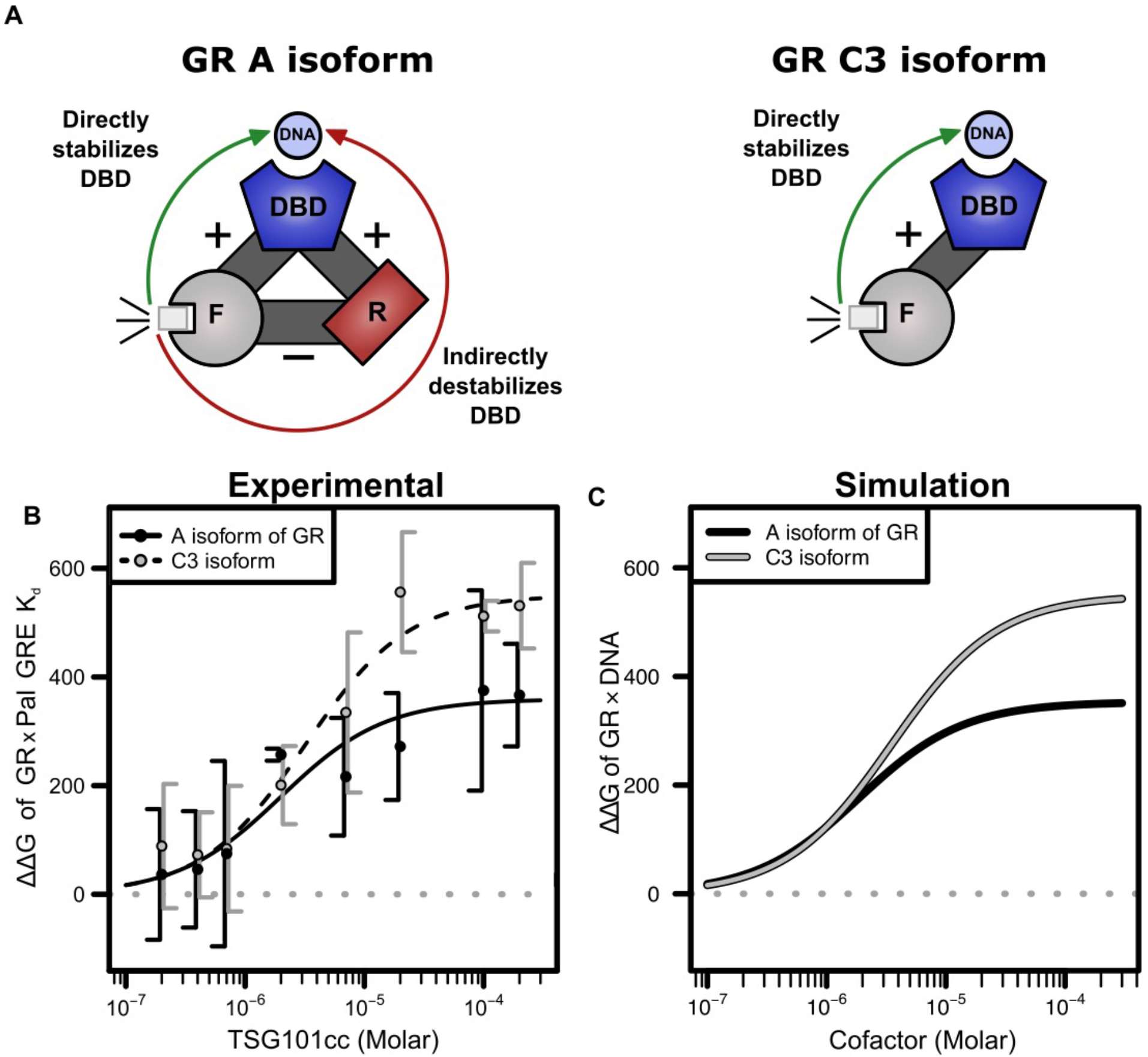
Energetic frustration in GR A is revealed by TSG101cc binding. **A)** Schematic representation of the interdomain thermodynamic coupling in the GR A NTD-DBD (left) and GR C3 NTD-DBD (right). The domains are labeled and portrayed as colored shapes connected by gray bars representing their thermodynamic coupling, either positive or negative. Binding of a cofactor (small gray box) to the GR F-domain can directly stabilize the DBD and promote binding of DNA via the F:DBD coupling; however, for GR A it will simultaneously destabilize the DBD and decrease DNA binding through its interaction with the R-domain of GR A. **B)** TSG101cc was titrated in multiple Pal GRE binding experiments, and each data set was fit individually, not globally. The ΔΔG of the y-axis was calculated as in Fig. 5. The lines of best fit are a simple binding equation: (ΔΔG_Max_ × TSG101cc ÷ K_d_) / (1 + [TSG101cc ÷ K_d_]). The points are the average of duplicate binding datasets ± one standard deviation, except at 100 μM TSG101cc five GR A and three GR C3 isoform datasets were used. Fitted values for TSG101cc K_d_ with GRE bound to GR: GR A K_d_ = 2.0 ± 2.9 μM (7.6 ± 0.5 kcal/mol), ΔΔG_Max_ = 360 ± 93 cal/mol; GR C3 K_d_ = 3.2 ± 2.3 μM (7.4 ± 0.3 kcal/mol), ΔΔG_Max_ = 550 ± 80 cal/mol (± 95% CI). **C)** A simulation of Part A using the EAM with values similar to our previous work (13). Attempts to directly fit our data proved the problem too unconstrained. See Supporting Information for explanation of the calculations here.

Importantly, the binding of TSG101cc to GR with and without DNA also illuminated the mechanistic impact of the regulatory R-domain that is present in the full-length GR A isoform, but which is missing in the GR C3 isoform. Specifically, thermodynamic analysis of the stability and DNA binding affinities of various isoforms of GR (13), revealed that the GR A isoform exists in a frustrated state, wherein the R-domain (Fig. 6A) decreases the probability of the GR A being in its active conformation (29). This occurs because the R-domain is both positively coupled and, therefore, stabilized by the DBD, and also negatively coupled, therefore destabilizing, to the F-domain, which is responsible for interaction with TSG101cc. Consequently, the impact of TSG101cc should have an overall greater impact on the GR C3 isoform, as the frustration present in the GR A isoform prevents the DBD from fully populating the high affinity state for DNA binding. In ensemble terms, under apparent saturating conditions of TSG101cc, only a subset of the ensemble is competent to bind DNA with high affinity. As such even though the DBD is identical in the GR A and GR C3 isoforms, the change in binding affinity with increasing TSG101cc saturates at a lower level for GR A NTD-DBD than it does for the GR C3 NTD-DBD (Fig. 6C).

### TSG101 co-localizes with GR in the nucleus, and both TSG101 and TSG101cc enhance transcription from glucocorticoid response elements (GREs)

To determine whether the *in vitro* effects reported above are consistent with the *in vivo* impact, GR-negative U2OS cells were transfected with TSG101 and GR. TSG101 was distributed throughout both the cytoplasm and the nucleus, in both endogenous and overexpressed conditions (Fig. 7 and Fig. S9). GR NTD-DBD localized to the nucleus, as is known to occur with GR lacking an LBD (13), and co-localized with both endogenous and overexpressed TSG101 in the nucleus (Fig. 7).

**Figure 7.**
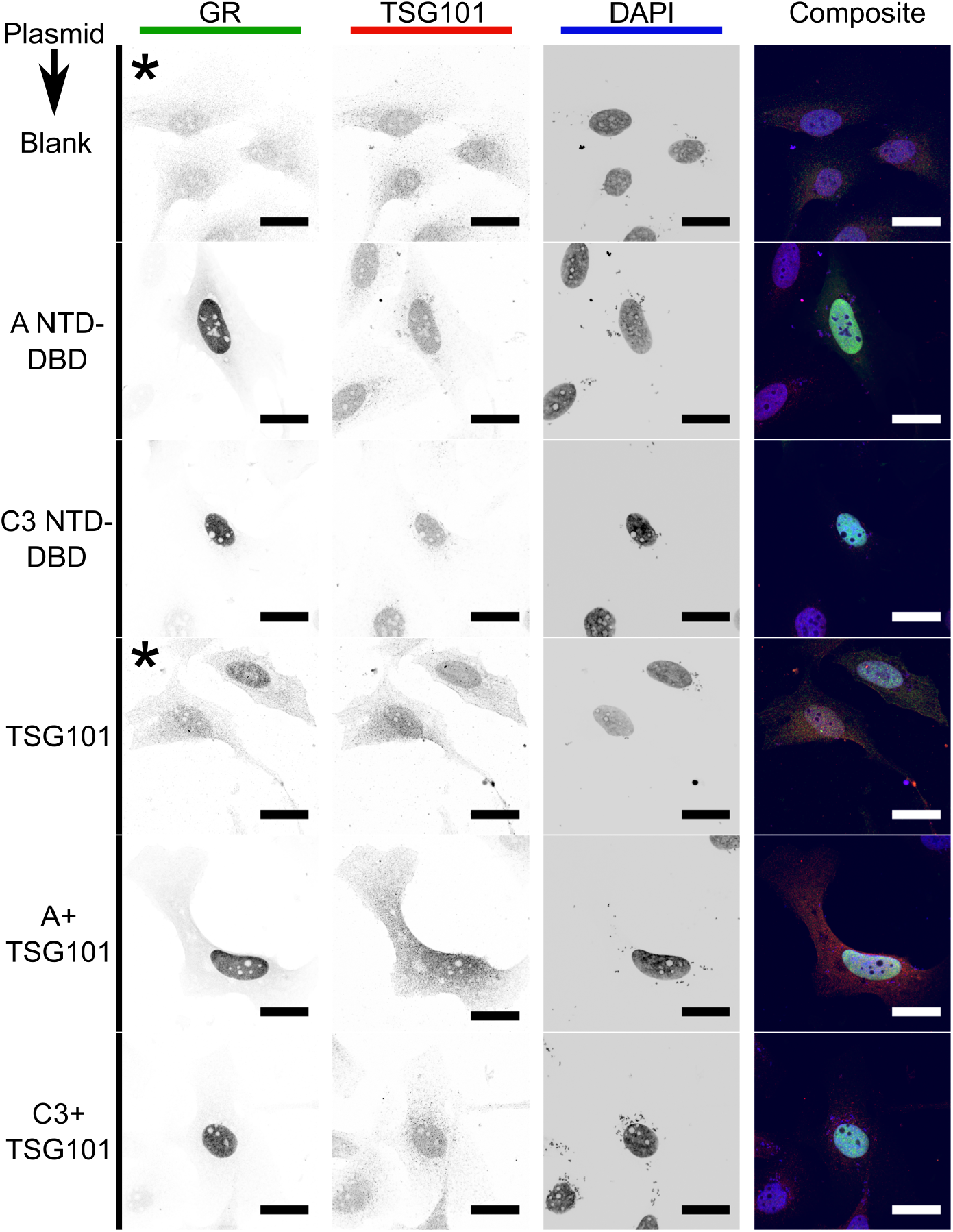
TSG101 and GR co-localize in U2OS nuclei. GR A/C3 NTD-DBD and TSG101 were detected by immunostaining. The expression plasmids transfected are indicated on the left of each image series. All scale bars are 25 μm. *In order to view the GR channel of cells without a GR expression plasmid, the microscope gain had to be increased about 1.4-fold. Examples shown are typical from six or more fields randomly examined (experimental groups) or three fields (blank).

Transfection of full-length TSG101 together with GR A or C3 NTD-DBD increased the expression of *Gaussia* luciferase driven by a GRE, in a dose-dependent manner (Fig. 8). The TSG101cc alone marginally increased GR transcriptional activity, but deletion of the TSG101 coiled-coil domain abrogated all stimulation (Fig. 8). Thus, TSG101 is clearly a co-regulator for GR gene induction, with the coiled-coil domain being both necessary and sufficient for TSG101-mediated stimulation of GR-driven transcription (although additional TSG101 regions appear to enhance the effect).

**Figure 8.**
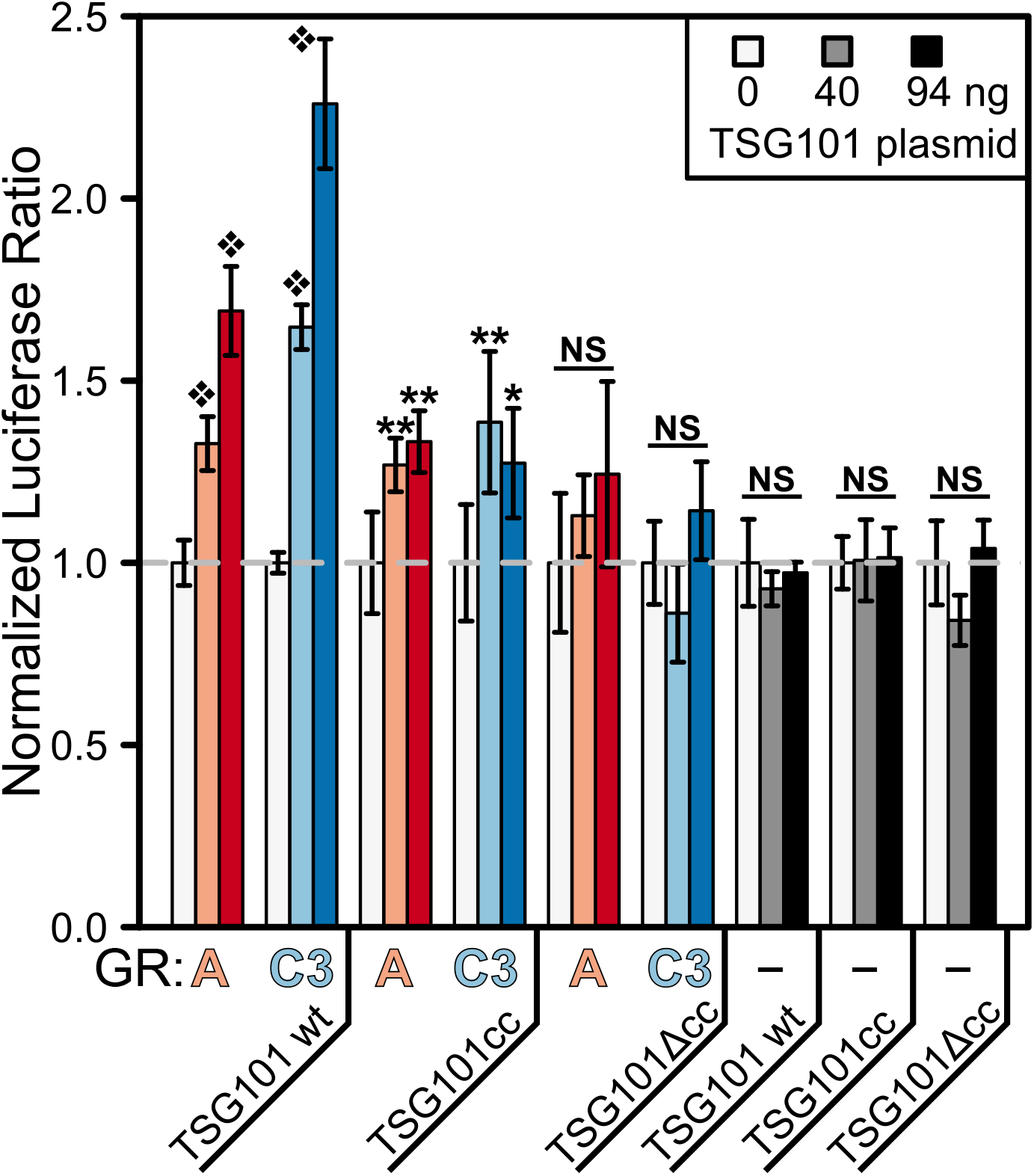
TSG101’s coiled-coil is necessary for enhancement of GR-driven transcriptional activation. Signal is from a reporter luciferase expressed from GRE promoter:luciferase DNA transfected into U2OS cells. For GR containing samples, a constant 6 ng of a given GR NTD-DBD plasmid was transfected. Error bars represent mean ± 95% CI of four samples. *P < 0.05, **P < 0.01, black diamond is P < 0.001. T-tests were one-sided with the alternative hypothesis being that the 0 ng group mean < experimental mean. Benjamini-Hochberg corrections were used throughout (57).

Finally, to test whether the increased activity was mediated through the AF1 region of the GR NTD, each of the eight translational isoforms of the GR were screened for activation by TSG101 in U2OS cells. Consistent with the effect being mediated by the AF1-containing region, the activity of all isoforms containing AF1 (i.e., GR-A, B, C1, C2, and C3) were enhanced by TSG101, while those without AF1 (i.e. GR-D1, D2, and D3) showed no response (Fig. S10). Further, when tested with full-length GR A (α), which contains the steroid binding LBD, TSG101 was able to hyper-activate GR so long as an activating steroid was administered (Fig. 9). Administration of a weak, mixed agonist (i.e., RU486) caused nuclear localization of GR (Fig. S11-12), but inhibited hyper-activation of GR by TSG101 and inhibited activation by dexamethasone (Fig. 9). These effects are likely related to RU486’s competition for the LBD’s steroid binding site and known recruitment of co-repressor proteins (30,31).

**Figure 9.**
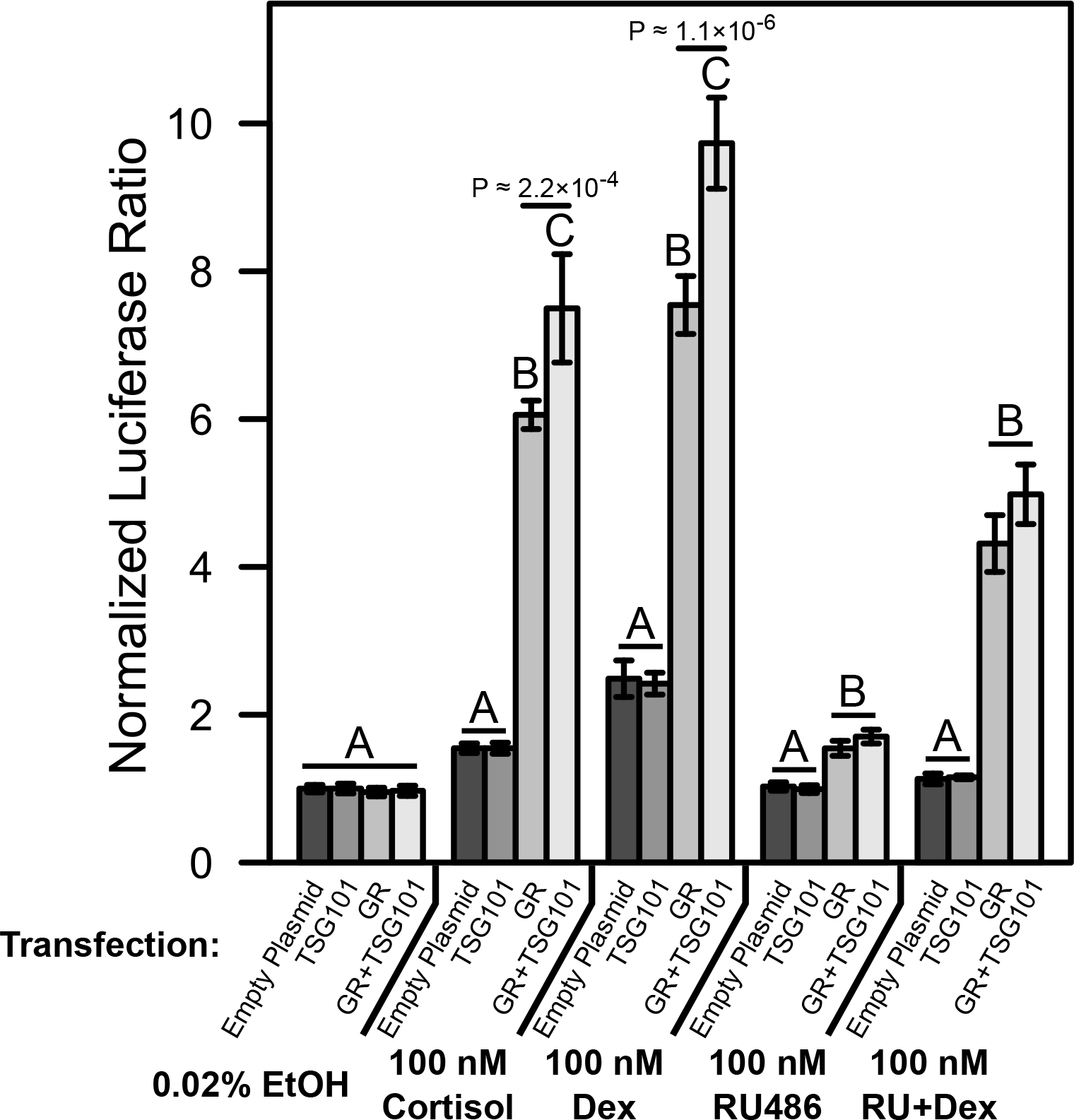
TSG101 enhances transcriptional activation driven by full-length GR A (α) Transcription of luciferase reporter from a GRE PAL-driven construct transfected into U2OS cells. Each data set was normalized to its respective empty plasmid transfection plus EtOH steroid vehicle; a higher ratio corresponds to more GR transcriptional activity. These example data were run in sextuplicate and are shown as means ± 95% confidence intervals. A Tukey HSD test was applied to each drug treatment, with at least a P < 0.01 defining different groups, labeled A, B, or C for each treatment. The actual P-values between B and C are listed in the figure. Dex = dexamethasone and RU+Dex = 100 nM of both RU486 and dexamethasone. RU486 is a partial agonist well known to partially block dexamethasone activation by binding to the steroid ligand binding site.

## Discussion

We have examined the allosteric behavior of ID-containing regulatory proteins by use of a well-studied ID transcription factor, GR, and the effect of TSG101 on its DNA binding affinity and activity. First, we confirm and extend earlier findings that the TSG101 coiled-coil domain binds the ID AF1-containing region of the GR (i.e. the F domain). We have improved the mapping of the interactions of TSG101cc and the GR NTD, and we provide novel evidence that binding between TSG101cc and GR causes tertiary structure to form in the NTD. Furthermore, we show that GR and TSG101 co-localize in the nucleus, supporting the view that they interact *in vivo.* Importantly, and counter to what was observed previously (21), TSG101 causes increased transcriptional activity of the GR. Hence, in the systems studied here, TSG101 is clearly a GR co-regulator that enhances transcription.

With regard to the functionally important regions of TSG101, we find that the coiled-coil domain is both necessary and sufficient for activation, although additional as yet undefined regions of TSG101 are needed for maximal effect. Toward this end, studies on the androgen receptor (AR) suggest that the TSG101 UEV and S-box domains may also be important for interactions with transcriptional machinery (25,32).

We also found that interaction of TSG101 with GR increases the affinity of the GR DBD for palindromic GRE DNA sequences, results that directly confirm and support the predictions of our model for allosteric proteins containing ID regions (16). Namely, ligand-induced binding and folding of an ID domain of a protein can allosterically influence the binding to a structured domain. Also consistent with our previous work, we found that the GR A isoform is energetically frustrated by its R-domain; thereby lowering the degree to which TSG101cc is able to promote GR A binding to DNA. The functional implications of these isoform dependent effects are currently not known.

In total, our results suggest a model whereby TSG101cc interacts with a helical region of GR AF1 (Fig. 10). Given the known functionally relevant binding partners unique to both TSG101 and GR in the cytosol, it is likely that TSG101 and GR interact weakly outside of the nucleus. However, it is clear that TSG101 can enter the nucleus and upon GR binding DNA, the interaction with TSG101 is greatly promoted (Fig. 10A). Importantly, our data are consistent with a model of GR and TSG101cc forming a dimeric triple helix, wherein the GR NTD folds back on the TSG101cc in a manner reminiscent of the anti-parallel ESCRT-I triple helical coiled-coil (20). Such a binding model would likely mimic the burial of hydrophobic surface that is seen in the heterotrimeric coiled-coil of the ESCRT-I complex, as shown previously (27). Confirmation of this hypothesis awaits further study.

**Figure 10.**
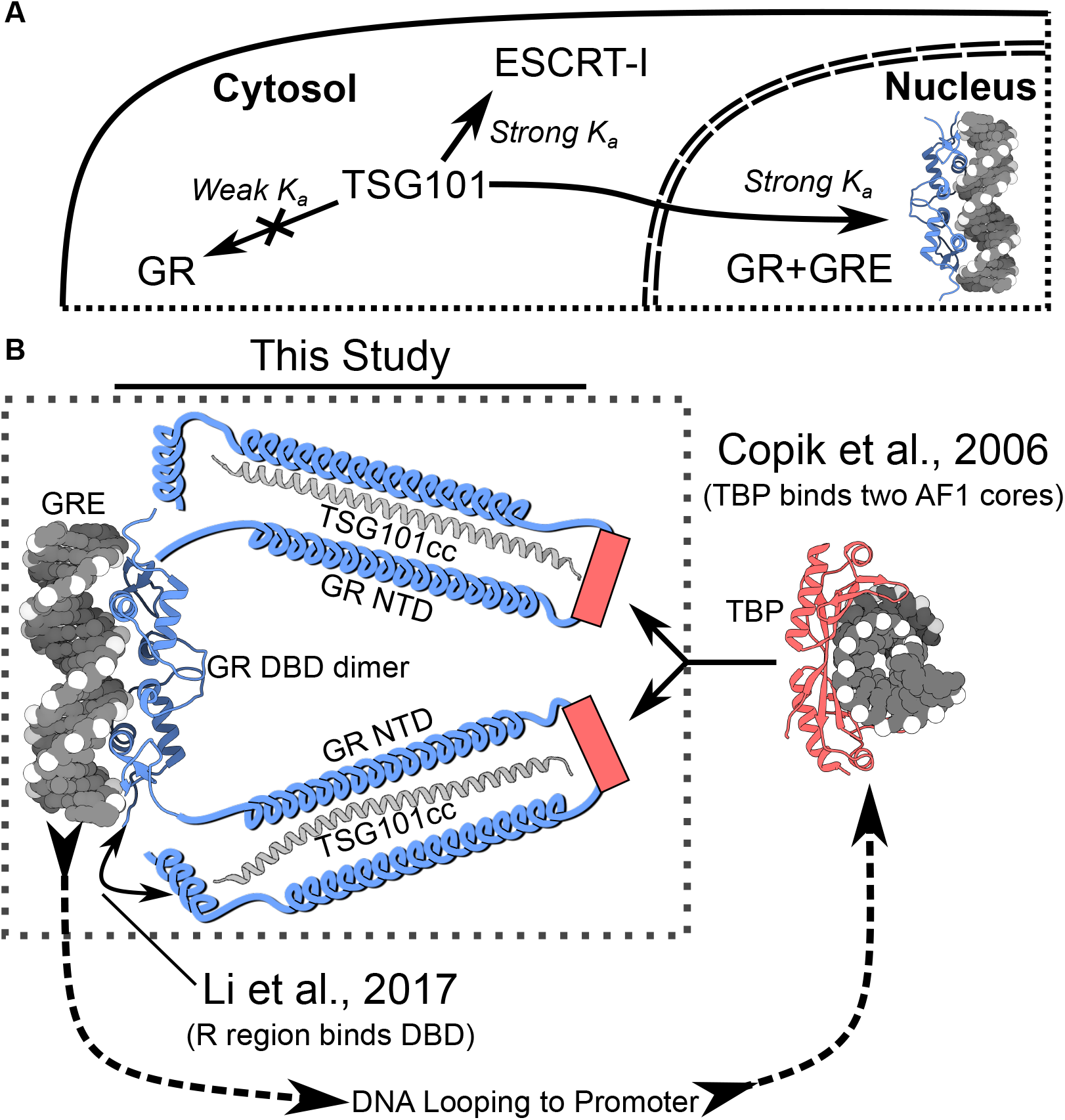
Model: TSG101cc in the nucleus directly binds to the GR NTD to enhance transcriptional activation. A model based on the results of this study is schematized in **A)** and the dash-lined boxed area of **B)**. **A)** A cell is partially sketched to illustrate that cytosolic TSG101 binds weakly to GR and better to other binding partners. Nuclear TSG101 binds GRE-bound GR more strongly than free GR. **B)** In the boxed area, GR (blue) bound to its GRE (grey) is shown. Each GR NTD forms a pseudo-trimer with one TSG101cc (grey coils, pdb:3iv1). The individual GR NTDs do not interact (our data only show heterodimers of GR NTD and TSG101cc). The R region of GR (N-term of the GR A NTD) interacts with the DBD (13), and this schematic model allows for that association. The data of Fig. S6 suggests that the R-region forms a helix, hence our depiction of R here. TSG101’s interaction with GR does not depend on the known TBP binding site (pink box), possibly leaving it open for interaction with TBP. On the **right**, TBP is shown in pink. Others demonstrated that TBP binds two AF1 core elements of GR (26). **Structures:** (33,58).

Finally, it is also important to note that the proposed model agrees with two other previously published reports (Fig. 10B). First, Copik, et al (26) found that the TATA box binding protein (TBP), which binds to the TATA box as a monomer (33), recruits and interacts with two different AF1 cores (Fig. 10B, right). Second, the anti-parallel alignment of the two regions of GR NTD that interact with TSG101cc yields a compelling and structurally reasonable model for how the R-region of GR, which is the most N-terminal region of the NTD, can interact with its own DBD, an interaction recently reported by Li, et al. (13) (Fig. 10B, left).

## Conclusion

We have shown that TSG101cc interacts with the GR NTD, confirming previous findings, and that the interaction enhances activity *in vivo*. Furthermore, we show that TSG101cc allosterically affects DBD binding of DNA and that DNA binding allosterically affects NTD binding of TSG101cc. Our results thus indicate that TSG101cc is a *bona fide* transcriptional co-regulator of GR function.

## Experimental Procedures

### Cloning of Plasmids Expressing Human GR or TSG101

All *E. coli* expression was achieved with the pJ411 plasmid of DNA2.0 (now ATUM, CA USA), and the cloning of most constructs was described previously (4,13). For the largest GR NTD constructs (A = a.a. 1—420, C3 = a.a. 99—420), the coding sequence was tagged with a 9x His tag on both the N and C-termini along with TEV cut sites to remove the tags. The smaller AF1 core of GR (a.a. 187—244) was tagged with an N-terminal 9xHis-TEV tag. The NTD-DBD constructs of GR (A = a.a. 1—525, C3 = a.a. 99—525) were tagged on just the N-terminus with a 6x His-Thrombin sequence (expression problems were observed with TEV or longer His tags). The open reading frame of the TSG101 coiled coil (a.a. 229—304) was tagged with a 9x His tag followed by a TEV site and a serine on the N-terminus (27). See Supporting Information for sequences used.

Besides luciferase, all U2OS expression of TSG101 or GR NTD-DBD was achieved with the pJ603 plasmid of DNA2.0, and the cloning of the GR constructs was described previously ((13); A = a.a. 1—525, B = a.a. 27—525, C1 = a.a. 86—525, C2 = a.a. 90—525, C3 = a.a. 99—525, D1 = a.a. 316—525). Note that the pJ603 plasmid uses a constitutive promoter (cytomegalovirus) to express the inserted gene. DNA2.0 generated the full-length TSG101 expression plasmid with codon optimization for human cell expression. See the Supporting Information for sub-cloning details. For studies of full-length GR, pEGFP GR was a gift from Alice Wong (Addgene plasmid #47504).

### Protein Expression and Purification

Expression and purification of TSG101cc and the various GR constructs used here has largely been described before (4,13,27) and detailed methods can be found in the Supporting Information text. See Figure S13 for examples of GR and TSG101 protein purity.

### Protease Protection, Cross-Linking, and Mass Spectrometry

Protease digestions were carried out in both the fluorescence anisotropy buffer (below) and 20 mM Na_2_HPO_4_ + 50 mM NaCl pH 7.2, with no observable differences. Saturation of GR required at least a thirty-fold excess of TSG101cc. To control for the total amount of protein present in each digest, the ratio of trypsin and targeted protein was held constant at 1000 μg protein:1 μg trypsin, except where noted. All digests occurred at room temperature after allowing the protein solutions at least 20 minutes to equilibrate. Each time point was taken by quenching an aliquot of reaction in Laemmli sample buffer and heating at 95° C for ~ 5 minutes.

Three cross-linkers were tested as follows, with their linker lengths in parentheses: BS3 (11.4 Å), DST (6.4 Å), and EDC + sulfo-NHS (0 Å) from Thermo-Fisher Scientific or ProteoChem. BS3 and EDC proved to be unusable in these circumstances, either because of too much or no cross-linking, respectively. The rest of this manuscript only refers to DST cross-linking. The DST cross-linking buffer was the same as the fluorescence anisotropy buffer (below). Initial reactions were carried out according to the manufacturer protocols, but because TSG101cc has a large number of lysines (11 out of 78 a.a.) there was an excessive amount of TSG101cc self cross-linking. After lowering the concentration of DST to about one-sixth or one-twelfth the recommended amount (0.5 mM used here) we were able to detect a faint GRxTSG101cc cross-link without TSG101cc background signal. See the Supporting Information for cross-linking and MALDI mass spectrometry details.

### Thermodynamic Fold Recognition

To gain insight into which structure(s) might be energetically compatible with the GR-NTD, the amino acid sequence (residues 1-420) was threaded against all domains contained in the Protein Data Bank (PDB) (34), as annotated in the *ECOD* database version 88 (35). The basic fold recognition algorithm used is described in (28), with the addition of three modifications specific to a database search using an IDP as query. First, thermodynamic environments were obtained using the *eScape* modification of *COREX* (36) instead of the original *COREX* algorithm (37), since the latter requires a crystal structure as input. Second, sequence:structure matches were scored using denatured state thermodynamic information in addition to native state information, as described in (38). Third, to increase sensitivity, only matches achieving reciprocally significant scores were considered, meaning that compatible environments between PDB and GR NTD sequence simultaneously exhibited compatible environments between GR NTD and PDB sequence. As described in (28), the significance (*p*-value) of a match was estimated by integrating a normal distribution between the limits of −∞ and the observed max[native + denatured] scores of the match, where the mean *μ* and standard deviation *σ* of the distribution were respectively defined as: *μ* = 0.00525 − 0.326\**L* and *σ* = 0.818*sqrt(*L*), where *L* is the length of the PDB domain.

To determine secondary structure, the structural coordinates of the aligned PDB hits were subsequently run through the DSSP algorithm built into Chimera (39,40). Of the entire PDB, 125 domains cleared the threshold for statistical significance (P < 10^−5^) and were used here. “Helix” described here is a combination of 3-10, π, and α-helix. The resulting per-residue secondary structure counts were then summed and normalized as a percentage of the total number of hits at a given residue. For presentation purposes, percentages were smoothed by a five amino acid rolling average with a step size of two.

### Fluorescence Anisotropy

The following buffer was used for all fluorescence anisotropy experiments shown here: 13.2 mM HEPES, 16 mM K Acetate, 67.2 mM NaCl, 4.2 mM MgCl_2_, and 0.84 mM TCEP all pH’d to 7.56 using 10 M KOH. It is difficult to purify large amounts of any GR construct that includes the disordered NTD. In order to maximize the high concentration end of our experiments, all the titrations presented here are dilution experiments: The fluorophore (pyrene-TSG101cc or 6FAM-DNA) was diluted into a concentrated solution of GR and the GR was then diluted out with a solution of fluorophore, with the concentration of fluorophore being kept the same throughout. The dilution solution was actively held at 20° C during the experiments to prevent thermal disequilibrium.

All experiments were at the very least duplicated with independent protein preparations, and all replicate data sets were fit globally to produce the presented values, except where noted. Equilibration times were varied to check for thermodynamic equilibrium, and because no obvious effects were discerned from 1 to ~ 30 minutes, most of the data presented were collected with 2 minute incubations between measurements. The only sensitive incubation step was the first measurement, which needed to be about 20 minutes long to establish thermal equilibrium. See Supporting Information for instrumental and fitting details.

### Cell Culture

U2-osteosarcoma cells (U2OS) were maintained in McCoy’s 5A medium, supplemented with 10% fetal bovine serum and antibiotics (streptomycin and penicillin). Cells were grown in 15 cm plates (Falcon) at 37° C in a 5% CO_2_ atmosphere and the medium was changed every 2– 3 days until passaging. Trypsin with EDTA was used for detaching cells during passaging. For transfection, cells were plated at a density of 3×10^4^ cells per well on 96-well plates or 3×10^5^ cells per well on 6-well plates and grown to ~ 90% confluence before transfecting. Xtreme GENE HP (Roche) was used to transfect DNA per the manufacturer directions with a resulting transfection efficiency of about 20%. For studies of full-length GR, cells were passaged using media supplemented with charcoal stripped FBS the day before transfection.

### Luciferase Assays

Most luciferase assays were done using 96-well plates (0.3 cm^2^ wells, Falcon), except comparison of TSG101 mutants proceeded by 6-well plates (9.4 cm^2^ wells, Sigma). The NEB *Gaussia* and *Cypridina* plasmids were used at 40 ng each per well for each 96-well assay (or ten times as much for 6-well plates). Two GR response elements were cloned into the promoter of the *Gaussia* plasmid and the *Cypridina* plasmid served as a control (13). GR expression plasmids were transfected at 0.6 ng per well and TSG101 plasmids were transfected from 0 up to 9.4 ng per well, with the empty pJ603 plasmid balancing the total transfection to 10 ng of pJ603 DNA (96-well plates).

For studies of full-length GR, 0.6 ng of eGFP-GR expression plasmid was co-transfected with either 9.4 ng of empty pJ603 plasmid or 8 ng of TSG101 pJ603 and 1.4 ng of empty pJ603. About 16—20 hours later, 1 mM hormone stocks in ethanol were diluted to 1.5 μM using media with charcoal stripped FBS. This hormone dilution or diluted ethanol control was then added immediately to the cells for a final hormone concentration of 100 nM and final ethanol concentration of 0.02%. In order to maintain the same concentration of ethanol across all experiments, when only one hormone was diluted to 1.5 μM, an equivalent volume of ethanol was added to yield the same final ethanol concentration as in dual-hormone experiments. Supernatants were collected about 24 hours after hormone addition.

*Gaussia* and *Cypridina* substrates were obtained from NEB or its original supplier, Targeting Systems. Either a Berthold or Promega luminometer were used for automated data collection.

### Confocal Microscopy

Cells were passaged onto German glass cover slips in 6-well plates. For the immunostaining data, there were two transfection procedures, both using 3 × 10^5^ cells per well. The first addressed whether or not TSG101 is in the nucleus of U2OS cells, with or without NTD-DBD GR, and was also used to determine full-length GR localization. Each pJ603 plasmid (TSG101 and/or GR) was transfected at 100 ng and sonicated salmon sperm DNA was used to bring the total amount of DNA to 1000 ng per well, and the cells were fixed about 40 hours after transfection. Transfection and induction of full-length GR proceeded as stated above.

The second procedure addressed how TSG101 localization changes in response to the luciferase assay conditions, over time. Transfection proceeded in the same manner as with luciferase assays. Cells were then collected for fixation 16 or 40 hours later. A cytosolic aberration appeared about 40 hours post-transfection in the group with the highest amount of TSG101 plasmid (Fig. S14-15). This coincided with cell death, as seen by floating cells in the associated wells. Overexpression of TSG101 previously caused both of these effects in literature data (41,42). Because we collected cells ~ 40 hours post transfection and both of our luciferases are constitutively excreted, our luciferase measurements are largely a time average of everything occurring before the shown artifact. We also observed repression of GR in some groups of older, high-passage cells, and the presented data are with low-passage cells only (< 7).

TSG101-tdTom was used as an alternative verification that TSG101 is in U2OS cell nuclei. For expression of TSG101-tdTom, 10^5^ cells were passaged and the expression plasmid was transfected at 15 ng per well and sonicated salmon sperm DNA was used to bring the total DNA amount per well to 500 ng. The TSG101-tdTom cells were fixed ~ 24 hours after transfection.

All immunostaining was similar to our previous work and details can be found in the Supporting Information (13).

All microscopy was done using the Integrated Imaging Center of Johns Hopkins University. The data shown were collected using either a Zeiss LSM 700 (immunostaining) or 780 (TSG101-tdTom) confocal microscope. DAPI was excited with a 405 nm diode laser, AlexaFluor 488 and eGFP (GR) were excited with a 488 nm laser, and both AlexaFluor 594 and tdTom (TSG101) were excited with a 561 nm laser. All imaging was done at room temperature with 400x or 630x magnification oil objectives (1.4 NA) using ZEN acquisition software. The gain and digital offset were optimized for the best signal-to-noise in each image; thus, the absolute intensities are not necessarily comparable. Analysis of images was done using the Fiji version of ImageJ (43,44).

## Acknowledgements

Professors Brand, Schroer, Hedgecock, Wendland, and Moudrianakis for sharing instruments, supplies, and advice. Dr Katie Tripp and the Center for Molecular Biophysics at Johns Hopkins University.

## Conflict of Interest

The authors declare that they have no conflicts of interest with the contents of this article.

## Author Contributions

JTW, JR, JOW, EBT, and VJH designed experiments. JTW, JR, MT, and JOW executed experiments. JTW and JOW analyzed data. All authors contributed to the final paper.

## Footnotes

This work was supported by an NIH training grant to the Cell, Molecular, Development, and Biophysics program in the biology department at JHU (T32-GM007231), and an NIH grant to VJH (R01-GM-126130). The content is solely the responsibility of the authors and does not necessarily represent the official views of the National Institutes of Health.

## The abbreviations used are

GR: glucocorticoid receptor
NTD: N-terminal domain
DBD: DNA-binding domain
LBD: ligand-binding domain
SHR: steroid hormone receptor
TSG101: tumor susceptibility gene-101
TSG101cc: TSG101 coiled-coil

